# Under pressure: phenotypic divergence and convergence associated with microhabitat adaptations in Triatominae

**DOI:** 10.1101/2020.07.28.224535

**Authors:** Fernando Abad-Franch, Fernando A. Monteiro, Márcio G. Pavan, James S. Patterson, M. Dolores Bargues, M. Ángeles Zuriaga, Marcelo Aguilar, Charles B. Beard, Santiago Mas-Coma, Michael A. Miles

## Abstract

**Background:** Triatomine bugs, the vectors of Chagas disease, associate with vertebrate hosts in highly diverse ecotopes. When these blood-sucking bugs adapt to new microhabitats, their phenotypes may change. Although understanding phenotypic variation is key to the study of adaptive evolution and central to phenotype-based taxonomy, the drivers of phenotypic change and diversity in triatomines remain poorly understood.

**Methods/Findings:** We combined a detailed phenotypic appraisal (including morphology and morphometrics) with mitochondrial *cytb* and nuclear ITS2 DNA-sequence analyses to study *Rhodnius ecuadoriensis* populations from across the species’ range. We found three major, naked-eye phenotypic variants. Southern-Andean bugs (SW Ecuador/NW Peru) from house and vertebrate-nest microhabitats are typical, light-colored, small bugs with short heads/wings. Northern-Andean bugs (W Ecuador wet-forest palms) are dark, large bugs with long heads/wings. Finally, northern-lowland bugs (coastal Ecuador dry-forest palms) are light-colored and medium-sized. Wing and (size-free) head shapes are similar across Ecuadorian populations, regardless of habitat or naked-eye phenotype, but distinct in Peruvian bugs. Bayesian phylogenetic and multispecies-coalescent DNA-sequence analyses strongly suggest that Ecuadorian and Peruvian populations are two independently-evolving lineages, with little within-lineage structuring/differentiation.

**Conclusions:** We report sharp naked-eye phenotypic divergence of genetically similar Ecuadorian *R. ecuadoriensis* (house/nest southern-Andean *vs*. palm-dwelling northern bugs; and palm-dwelling Andean *vs*. lowland); and sharp naked-eye phenotypic similarity of typical, yet genetically distinct, southern-Andean bugs from house and nest (but not palm) microhabitats (SW Ecuador *vs*. NW Peru). This remarkable phenotypic diversity within a single nominal species likely stems from microhabitat adaptations possibly involving predator-driven selective pressure (yielding substrate-matching camouflage coloration) and a shift from palm-crown to vertebrate-nest microhabitats (yielding smaller bodies and shorter heads and wings). These findings shed new light on the origins of phenotypic diversity in triatomines, warn against excess reliance on phenotype-based triatomine-bug taxonomy, and confirm the Triatominae as an informative model-system for the study of phenotypic change under ecological pressure.

**Author summary:** Triatomine bugs feed on the blood of vertebrates including humans and transmit the parasite that causes Chagas disease. The bugs, of which 150+ species are known, are highly diverse in size, shape, and color. Some species look so similar that they are commonly confused, whereas a few same-species populations look so different that they were thought to be separate species. Despite the crucial role of naked-eye phenotypes in triatomine-bug identification and classification (which are essential for vector control-surveillance), the origins of this variation remain unclear. Here, we describe a striking case of phenotypic divergence, with genetically similar bugs looking very different from one another, and phenotypic convergence, with bugs from two genetically distinct populations (likely on their way to speciation) looking very similar – and all within a single nominal species, *Rhodnius ecuadoriensis*. Phenotypically divergent populations occupy different ecological regions (wet *vs*. dry) and microhabitats (palm-crowns *vs*. vertebrate nests), whereas convergent populations occupy man-made and nest (but not palm) microhabitats. These findings suggest that triatomines can ‘respond’ to ecological novelty by changing their external, naked-eye phenotypes as they adapt to new microhabitats. We therefore warn that phenotypic traits such as overall size or color may confound triatomine-bug species identification and classification.

## Introduction

Triatomine bugs transmit *Trypanosoma cruzi* among the mammalian hosts they associate with in shared microhabitats [1,2]. Bugs that occur in man-made habitats may transmit the parasite to humans, fueling the spread of Chagas disease [1,3]. When these blood-sucking bugs adapt to new habitats, their phenotypes may change [4]. Over the last few decades, molecular studies have identified examples of phenotypic convergence or divergence at several systematic levels [5–9]. Perhaps most remarkably, molecular phylogenetic analyses suggest that none of the three genera where the main Chagas disease vectors belong (*Triatoma, Panstrongylus*, and *Rhodnius*), which are all defined after morphological characters [1], is monophyletic [8–14]. Some named species have been shown to be phenotypic variants of another species [8]; for example, *Triatoma melanosoma* is now regarded as a *T. infestans* chromatic variant [15,16]. Conversely, some named species have been shown to include several cryptic taxa [8]; for example, *Rhodnius robustus* (*s.l.*) is composed of at least five distinct lineages, two of which have been formally described as *R. montenegrensis* and *R. marabaensis* [8,17–21].

The sometimes striking variation of triatomine-bug phenotypes has been attributed to a propensity of morphological characters to change in response to shifting habitat features [2,4,22]. Thus, within-species divergence may be driven by habitat shifts (e.g., wild to domestic) involving subsets of genetically homogeneous populations [4], and the use of similar habitats by genetically distinct populations may result in convergence or in the retention of ancestral phenotypes [2,23]. When using morphological characters only, therefore, taxonomists are in peril of describing spurious species or overlooking cryptic taxa [8]. Despite the practical importance of accurate taxonomic judgment when the organisms of interest transmit a life-threatening parasite, the degree, direction, and underlying causes of phenotypic change and diversity in triatomines remain obscure.

In this paper, we combine a detailed phenotypic characterization, qualitative and quantitative, with mitochondrial (mt) and nuclear (n) DNA sequence analyses to study *Rhodnius ecuadoriensis* populations spanning most of the geographic/ecological range of the species (Fig 1). *Rhodnius ecuadoriensis* is a major domestic vector of *T. cruzi* in western Ecuador and northwestern Peru [24–26]. In Ecuador, northern wild populations are primarily associated with the endemic *Phytelephas aequatorialis* palm in both Andean wet forests and lowland dry forests; in the dry inter-Andean valleys of southwestern Ecuador, where palms are rare or absent, wild *R. ecuadoriensis* seem to primarily associate with arboreal squirrel nests [1,2,27–33]. The natural habitats of Peruvian populations remain largely unknown, with a few records suggesting association with vertebrate nests/refuges in hollow trees and perhaps cacti [2,26,27,34]. In addition, some *R. ecuadoriensis* populations have adapted to live in and around houses in coastal Ecuador and, especially, in the dry valleys of southwestern Ecuador and northwestern Peru – where the bugs contribute to endemic Chagas disease [24–26,35–39]. Our comparative phenotypic and genetic analyses reveal that phenotypic divergence and convergence can both occur within a single nominal triatomine-bug species, and suggest that microhabitat adaptations likely play a crucial role in this phenomenon.

**Fig 1.**
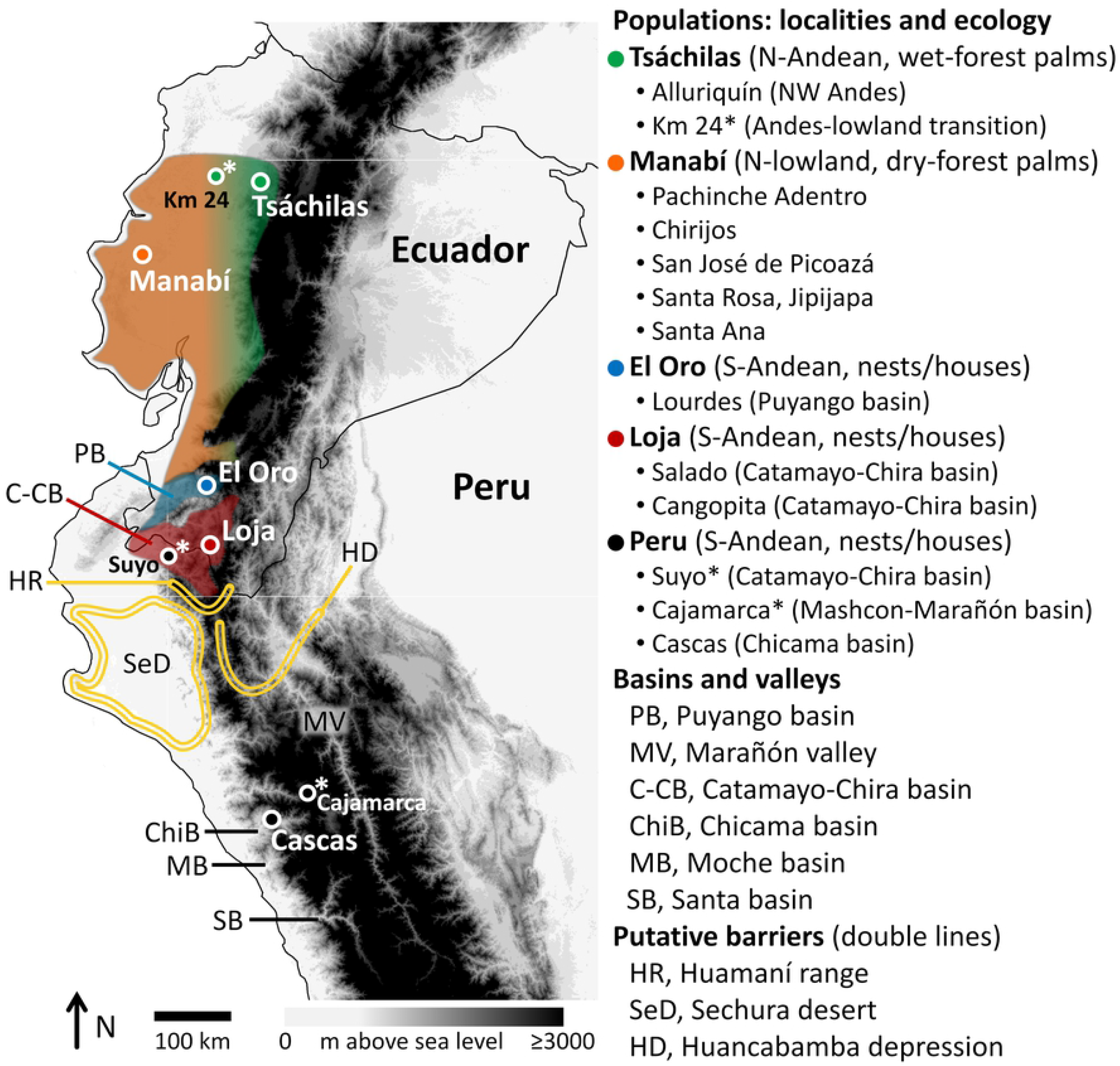
Sampling of *Rhodnius ecuadoriensis* populations. The map shows the approximate known distribution of *R. ecuadoriensis* in Ecuador, including the western Andean wet forests (green shade), the drier coastal lowlands (orange shade), and the southern, dry inter-Andean valleys associated with the Puyango (PB, blue shade) and Catamayo-Chira basins (C-CB, red shade). The approximate geographic location of each study population in Santo Domingo de los Tsáchilas, Manabí, El Oro, Loja, and Peru is indicated by, respectively, green, orange, blue, red, and black circles; just one specimen was available from sites with asterisks (Km 24, Suyo, and Cajamarca). Most Peruvian material came from Cascas, in the middle-upper Chicama basin (ChiB), La Libertad. Putative barriers to past or current bug dispersal are indicated with gold-colored double lines: the Huamaní range (HR), which closes the C-CB to the south; the Sechura desert (SeD) on the northern Peruvian coastal plains; and the Huancabamba depression (HD). The semiarid Santa river basin (SB) appears to mark the southern limit of *R. ecuadoriensis*’ range. *Rhodnius ecuadoriensis* also occurs along the middle-upper (inter-Andean) stretches of the Marañón river valley (MV). See details in S1 Table and main text.

## Methods

### Origins of bugs and qualitative phenotype assessment

We compared *R. ecuadoriensis* type specimens (Laboratório Nacional e Internacional de Referência em Taxonomia de Triatomíneos [LNIRTT], Fiocruz, Brazil; [40]) and the description of the type material [41] with field-collected Ecuadorian bugs including (i) southern-Andean bugs caught in/around houses of El Oro (Puyango river basin) and Loja (Catamayo-Chira basin); and (ii) northern bugs from *Ph. aequatorialis* palms of Manabí (lowland dry forest) and Santo Domingo de los Tsáchilas (Andean wet forest; ‘Tsáchilas’ hereafter) (Fig 1). CAC Cuba (University of Brasília, Brazil) supplied additional, field-caught southern-Andean Peruvian bugs from dwellings of Suyo (department of Piura, Catamayo-Chira basin) and Cascas (department of La Libertad, Chicama basin) [39] (Fig 1). Finally, bugs from two colonies founded with material collected from, respectively, *Ph. aequatorialis* palms of Manabí and houses of northwestern Peru (department of Cajamarca, Mashcon-Marañón basin) were supplied by J Jurberg (LNIRTT) (S1 Table). We conducted a detailed review of external morphological and chromatic characters central to classical triatomine-bug taxonomy [1,41], and placed the results in the broader context of what we know about the systematics, biogeography, and ecology of *R. ecuadoriensis* [1,2,8,18,25–36,38,39]. In particular, we emphasize that northern populations exploit palm-crown microhabitats just like most *Rhodnius* species do [2,29], whereas wild southern-Andean populations are associated with vertebrate tree-nests in dry ecoregions where palms are either rare or absent [2,8,25–34]. Our sampling thus captures this key ecological difference, with northern Tsáchilas and Manabí bugs representing primarily palm-dwelling populations and southern El Oro, Loja, and Peru bugs representing primarily nest-dwelling populations.

### Traditional morphometrics – heads

We used 79 adult *R. ecuadoriensis* specimens for this part of the study; specimen details are presented in S1 Table. We measured lateral- and dorsal-view, calibrated head images (Fig 2) and calculated descriptive statistics for the whole sample and for ecological (palm *vs*. house/nest populations) and geographic groups. To assess head-size variation, we estimated population means (over all head measurements) and likelihood-profile 95% confidence intervals (CIs) by fitting Gaussian generalized linear models (identity link-function, no intercept) using package *lme4* 1.1-21 [42] in R 3.6.3 [43]. For multivariate analyses, log-transformed data were centered by row to remove isometric size; the resulting ‘log-shape ratios’ [44,45] were used as input for principal component analysis (PCA) on covariances. The derived principal components (PCs or ‘shape variables’) were submitted to canonical variate analysis (CVA). We assessed the overall significance of multivariate CVA using Wilks’ λ statistic [46]. We computed canonical vectors (CVs) and used the first two CVs to plot the position of each specimen on the shape discriminant space; ‘convex hulls’ enclosing all points within each group were overlaid on the plots. Finally, a space of size-free shape variables was constructed by explicitly removing size (represented by PC1) from the measurements; for this, residuals of linear regression of PC1 on each measurement were used as new variables for size-free CVA [47–49]. The derived CV1 and CV2 were plotted as above. Different parts of these analyses were conducted using R 3.6.3 [43], JMP 9.0 (SAS Institute, Cary, NC), and NTSYS 2.10y [50].

**Fig 2.**
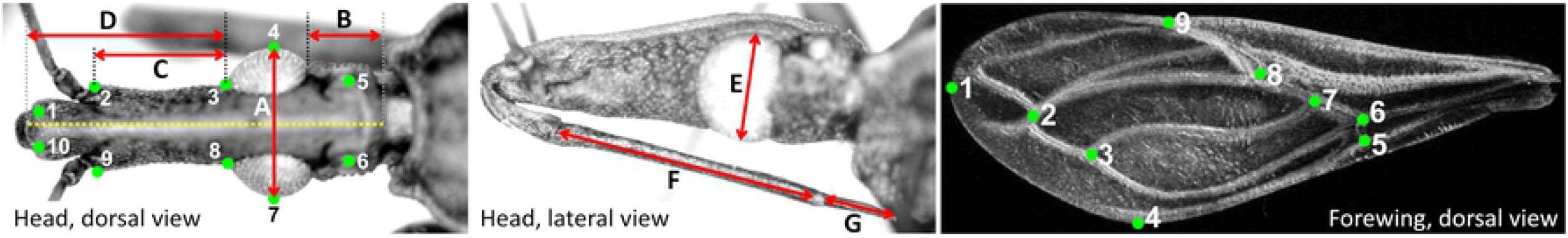
Measurements and landmarks used in morphometric analyses. Head measurements used for traditional morphometrics: A, maximum width across the eyes; B, postocular distance (posterior eye limit to head/neck limit); C, length of antenniferous tubercle (anterior eye limit to distal tip of tubercle); D, anteocular distance (anterior eye limit to base of anteclypeus); E, maximum diameter of the eye; F, length of second rostral segment; and G, length of third rostral segment. The yellow dotted line indicates head length, which we used, together with A, to compute head length:width ratios. Green dots show the landmarks used for geometric morphometrics.

### Geometric morphometrics – heads and forewings

Dorsal-head and forewing images of, respectively, 84 and 82 adult bugs (details in S1 Table) were digitized for a series of two-dimensional coordinates (Fig 2) using *tpsDig* 1.18 [51]. Raw coordinates were subjected to the Procrustes superimposition algorithm [52] and thin plate spline (TPS) analysis using *tpsRelw* 1.18 [53]. We used TPS to compute ‘partial warps’ with affine (global stretching) and non-affine (non-linear localized distortions or ‘shape changes’) components. PCA of partial warps yielded shape components (‘relative warps’), which were subjected to CVA as described above.

### Molecular analyses

We used 72 *R. ecuadoriensis* specimens for mtDNA analyses and a subset of 17 bugs for nDNA analyses (details in S1 Table); *R. colombiensis*, *R. pallescens*, and *R. pictipes* were used as outgroup taxa. We extracted DNA from bug legs using DNeasy kits (Qiagen, Valencia, CA). A 663 base-pair (bp) fragment of the mitochondrial cytochrome *b* gene (*cytb*) and a 707–715 bp fragment of the nuclear ribosomal second internal transcribed spacer (ITS2) were amplified, purified, and Sanger-sequenced as previously described [9,11,17]. We visually inspected the chromatograms of forward and reverse DNA strands with SeqMan Lasergene 7.0 (DNASTAR Inc., Madison, Wisconsin); in inspecting ITS2 chromatograms, we particularly checked for the ‘double signal’ typical of paralogous pseudogene sequences [54,55]. We aligned consensus and outgroup sequences in MAFFT 7.0 [56], using the L-INS-i algorithm and further manual fine-tuning. We then computed descriptive statistics using MEGA X [57].

The best-fitting model of base substitution for each marker was selected using the Bayesian information criterion (BIC) in bModelTest 1.2 [58]. We used the *BEAST 0.15 package of the BEAST 2.6 platform [59,60] to reconstruct Bayesian locus-specific phylogenetic trees and multispecies-coalescent species trees [59], with three independent runs (5×10^7^ generations) for each analysis. For locus-specific trees, we used the coalescent model and sampled parameters every 10,000 generations; for species trees, we used the Yule model of speciation and sampled parameters every 50,000 generations. We evaluated (using Tracer 1.7 [61]) parameter convergence and proper mixing by inspecting individual chains and by checking that effective sample sizes (ESSs) were sufficiently large; in our case, all ESSs were ≥10^4^.

It has been suggested, based on limited mtDNA data, that *R. ecuadoriensis* may comprise two distinct lineages – one primarily from Ecuador (‘group I’) and the other primarily from Peru (‘group II’) [18]. We set to formally assess the data support for this ‘two-lineage’ hypothesis (H_1_), relative to the null hypothesis of a single lineage (H_0_), by computing and comparing the marginal likelihood (mL) and posterior probability of each hypothesis [62,63]. To do this, we first used our *cytb* and ITS2 sequence data to estimate species trees under both H_1_ (with each *R. ecuadoriensis* sequence assigned to a pre-defined, geography-based group – ‘Ecuador’ or ‘Peru’; Fig 1) and H_0_ (with all *R. ecuadoriensis* sequences assigned to a single group). We then estimated each species tree’s mL with two approaches: (i) nested sampling [63,64] as implemented in the NS 1.1 package [64] of BEAST 2.6 [60], with 5×10^6^ MCMC generations, 2×10^4^ sub-chain length, and five active points; and (ii) path sampling (also known as ‘thermodynamic integration’) [63,65] as implemented in the Model Selection 1.0.1 package of BEAST 2.6 [60], with a pre-burn-in of 2×10^5^ MCMC iterations followed by 90 steps of 2×10^6^ iterations (50% burn-in) and each step repeated 500 times. Using the hypothesis-specific log-mL values, we finally computed log-Bayes factors as log-BF = log-mL(H_1_)-log-mL(H_0_) [62]. For two hypotheses with equal prior probabilities, Pr(H_1_) = Pr(H_0_) = 0.5, the Bayesian posterior probability (given the data, D) of H_1_ is Pr(H_1_|D) = BF/(1+BF), and the posterior probability of H_0_ is therefore Pr(H_0_|D) = 1-Pr(H_1_|D). Kass and Raftery [62] proposed a set of rules of thumb, derived from those first suggested by Jeffreys [66], to grade the evidence in favor of H_1_: the evidence is weak when 1 ≤ BF < 3; positive when 3 ≤ BF < 20; strong when 20 ≤ BF < 150; and very strong when BF ≥ 150. In a two-hypotheses context like ours, this means that evidence in favor of H_1_ (or against H_0_) would be deemed ‘very strong’ only if examination of the data changed the 1:1 prior odds (Pr(H_1_) = Pr(H_0_) = 0.5) to a posterior odds of at least 150:1, so that Pr(H_1_|D) ≥ 0.993 and Pr(H_0_|D) ≤ 0.007.

We note that the *R. ecuadoriensis cytb* sequences studied by Villacis et al. [67,68] are not available in NCBI’s GenBank or in EMBL-EBI’s European Nucleotide Archive (ENA). We also note that there are five *R. ecuadoriensis cytb* sequences in GenBank. Sequence AF045715 [69] is only 399 bp long, and sequences KC543508–KC543510 [70] are identical to some of the haplotypes we found (details below). Finally, KC543507 from a Manabí bug [70] differs at three positions from two of our Manabí haplotypes; one of those substitutions, however, yields a thymine at a second codon position where cytosine (position 233 in S1 Alignment) is conserved across all *cytb* sequences from *Rhodnius* species we have been able to examine (see, e.g., [8]). This singular non-synonymous substitution (valine instead of alanine as in all other *Rhodnius* except *R. prolixus* and some *R. robustus*, which have threonine) suggests that KC543507 [70] may contain base-call errors, and we therefore excluded this particular sequence from further consideration. Our analyses, in sum, are based on all reliable sequences of reasonable length (here, 663 bp) that, as far as we know, are currently available for the two loci we investigated.

## Results

### Qualitative phenotypic assessment

All southern-Andean bugs from El Oro, Loja, and Peru had comparable, typical phenotypes [1,41], whereas northern-Andean specimens from wet-forest palms (Tsáchilas) were very large and dark bugs with long, slender heads; northern-lowland bugs from dry-forest palms (Manabí) had intermediate phenotypes (Figs 3 and 4). Below and in Table 1 we present summary descriptions of these diverse putative *R. ecuadoriensis* phenotypes (or ‘forms’); detailed descriptions are provided in S1 Text.

**Fig 3.**
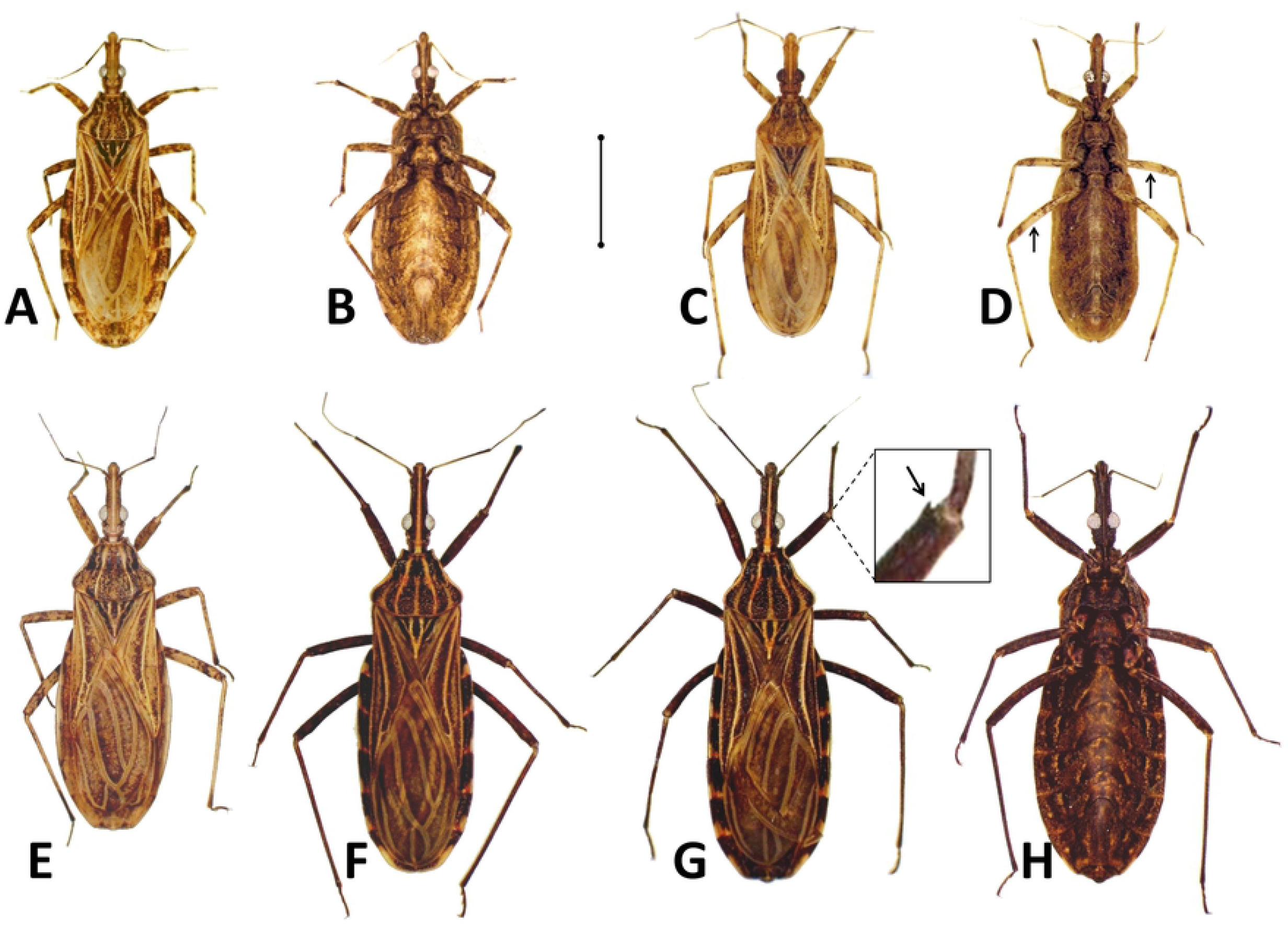
Phenotypic diversity in *Rhodnius ecuadoriensis*. **A–D**, southern-Andean house/nest populations: Loja female, dorsal (**A**) and ventral (**B**) views; and Peru male, dorsal (**C**) and ventral (**D**) views; the arrows in **D** indicate the lighter central area of femora (see text). **E–H**, northern, palm-dwelling populations: northern-lowland Manabí female, dorsal view (**E**); northern-Andean Tsáchilas male, dorsal view (**F**); northern-Andean Tsáchilas female, dorsal view (**G**; the inset highlights the well-developed denticle in the distal tip of the fore femora); and northern-Andean Tsáchilas female, ventral view (**H**). Scale bar, approx. 5 mm.

**Fig 4.**
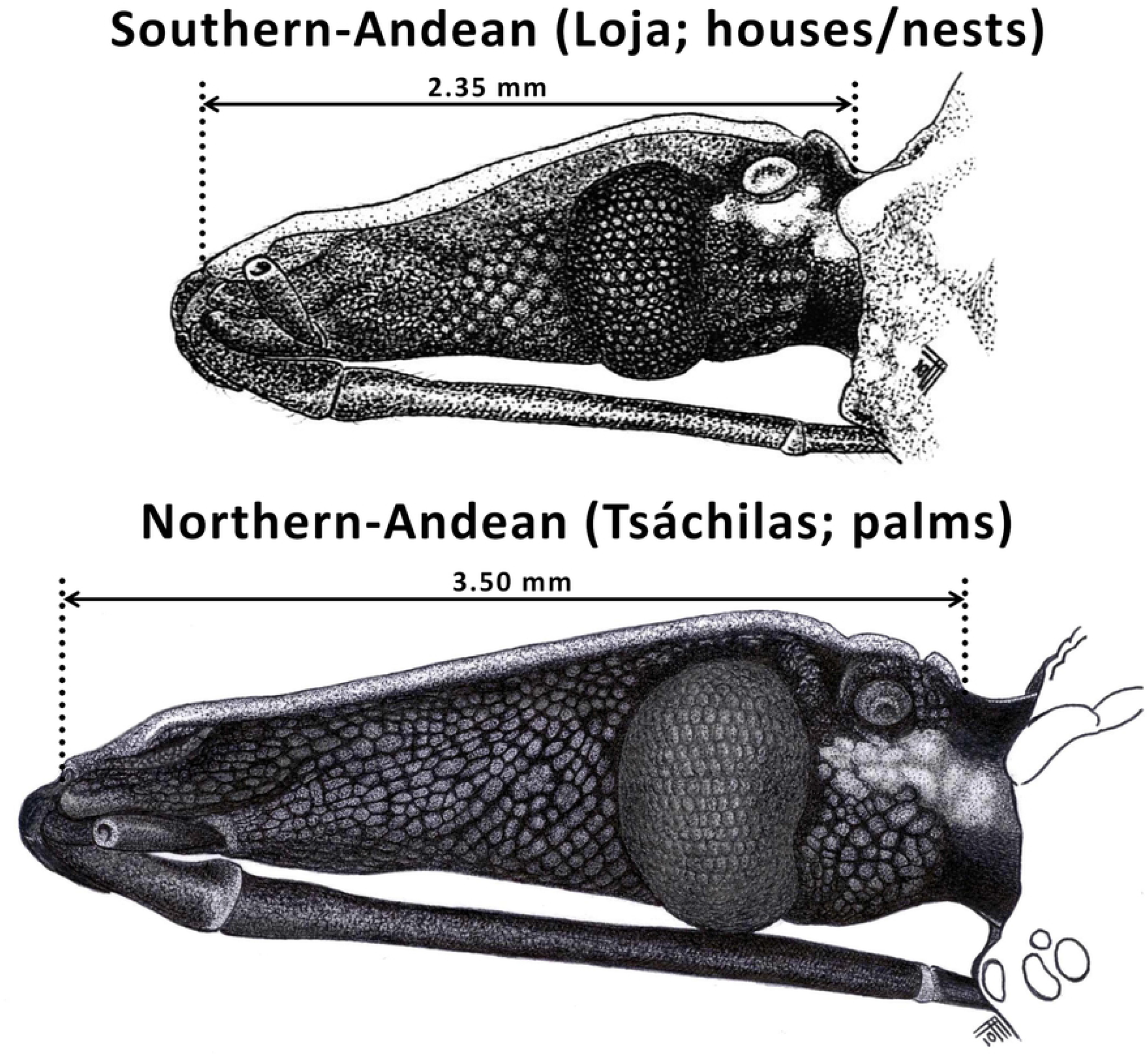
Heads (lateral view) of southern-Andean and northern-Andean *Rhodnius ecuadoriensis*: typical house/nest bugs (Loja) *vs*. atypical Andean palm bugs (Tsáchilas). Note the striking divergence, which clearly falls within the range of what are normally considered interspecies differences in the Triatominae [1]. Drawings by FA-F.

**Table 1.** Summary of phenotypic and ecological variation in *Rhodnius ecuadoriensis*.

| Form | Phenotypic traits<sup>a</sup> | Known ecology<sup>b</sup> |
| --- | --- | --- |
| Typical<sup>c</sup> (southern-Andean; houses/nests) | Similar to species types; pale yellowish; mottled pattern conspicuous; small size (body length ~13–14 mm); short and stout heads; MPP pointed | Squirrel and bird nests, hollow trees with *Didelphis*, houses and peridomestic structures (mainly hen nests, dovecotes, and guinea-pig pens); inter-Andean valleys with dry forests in southwestern Ecuador and northwestern Peru |
| Manabí (northern-lowland; dry-forest palms) | Light brown-yellowish; mottled pattern conspicuous; intermediate size (~15–16 mm); heads longer than in typical specimens; MPP pointed | *Phytelephas aequatorialis* palms plus, occasionally, bird, squirrel, rat, or mouse nests and man-made habitats; lowland dry forests in western Ecuador |
| Tsáchilas (northern-Andean; wet-forest palms) | Dark brown-black with brown-reddish markings; mottled pattern inconspicuous due to dark background color; large (~17–18 mm); long and narrow heads; MPP generally truncated | *Phytelephas aequatorialis* palms; Andean wet premontane forests (~300–1800 m above sea level) in western Ecuador |

MPP, median process of the pygophore (see S1 Fig)
<sup>a</sup> Detailed descriptions are provided in S1Text
<sup>b</sup> Refs. [1,2,24–39]
<sup>c</sup> Using the original description of the species [41] as the benchmark; see also ref. [1]

#### Southern-Andean house/nest forms

Bugs caught in/around houses in southwestern Ecuador (El Oro, Puyango basin; and Loja, Catamayo-Chira basin) were virtually identical to the type material – very small triatomines, light brown-yellowish with dark brown stripes and irregular markings on the body and appendages [1,41] (Fig 3). Peruvian bugs were collected in/around houses of the dry middle-upper Chicama basin (Cascas, La Libertad, ~350 km south of our fieldwork sites in Loja), except for one specimen collected in Suyo, Piura, within the Catamayo-Chira basin (Fig 1, S1 Table). The overall aspect of Peruvian bugs largely matches that of type material (Fig 3). However, Chicama-basin bugs are noticeably lighter-colored than southern-Andean Ecuadorian bugs; this is more evident on the legs, where the dark mottled pattern is limited to small clusters of dots and stripes on the basal and distal thirds of femora and tibiae (Fig 3). The posterior lobe of the pronotum is also lighter than in Ecuadorian material, and Chicama-basin bugs are more slender than the typical specimens from El Oro-Loja. Similar to type material, the heads of these Peruvian specimens are noticeably short and stout (Fig 3; see also S1 Text).

#### Northern-lowland palm forms

*Phytelephas aequatorialis* palms often harbor *R. ecuadoriensis* populations in the central coastal province of Manabí, Ecuador [2,28,29,31]. The overall aspect and coloration of these lowland palm-dwelling bugs largely matches those of typical *R. ecuadoriensis*, but Manabí bugs have larger bodies and longer, more slender heads (Fig 3; see also S1 Text).

#### Northern-Andean palm forms

In 1998, a male *Rhodnius* specimen was collected at light near Alluriquín, Santo Domingo de los Tsáchilas, ~900 m above sea level on the central-western Ecuadorian Andes foothills (Fig 1). This wet premontane-forest site is within the range of *R. ecuadoriensis*, which is not shared by any other known *Rhodnius* species [8,18,25,71], but the morphology and coloration of the specimen differed strikingly from those of the type material (Figs 3 and 4). Field surveys in Alluriquín yielded abundant material from *Ph. aequatorialis* palms [28,72]. These bugs are much larger and darker, and have much longer heads, than typical *R. ecuadoriensis*, but are smaller than the closely-related, light-colored *R. pallescens* and *R. colombiensis* [1,73] (Figs 3, 4, and S2 Fig). Naked-eye phenotype differences between these northern-Andean bugs and typical southern-Andean house/nest specimens are in the range customarily associated with distinct species in the Triatominae – and tend towards the ‘highly divergent’ extreme of that range if we consider closely-related species within the genus *Rhodnius* [1,5,8,17–19] (Table 1, Figs 3 and 4; S1 Text).

#### Traditional morphometrics – heads

Northern-Andean bugs from Tsáchilas palms clearly had the largest heads in our sample; northern-lowland bugs from Manabí palms were smaller than Tsáchilas specimens but larger than southern-Andean bugs – among which those from Loja had the smallest heads and those from Peru were larger on average (Fig 5A). Overall, the heads of northern, palm-dwelling bugs were not only larger (Fig 5B) but also more elongated (Fig 5C) than those of southern-Andean house/nest bugs. CVA revealed large among-group differences (Wilks’ λ = 0.024, *p* < 0.0001), with a negative correlation between CV1 scores and head size (Fig 5D). Discrimination between northern palm-dwelling and southern-Andean house/nest populations was complete. Some overlapping occurred between El Oro and Loja, and a single Peruvian specimen was firmly nested within the Loja cluster (asterisk in Fig 6A). This specimen was collected in Suyo, ~20–50 km from our fieldwork sites in Loja and also within the Catamayo-Chira basin (Fig 1). When size effects were explicitly removed (see **Methods**), all Ecuadorian populations (plus the Suyo bug) were remarkably similar to one another, with bugs from Peru appearing as the most distinct (Fig 5E).

**Fig 5.**
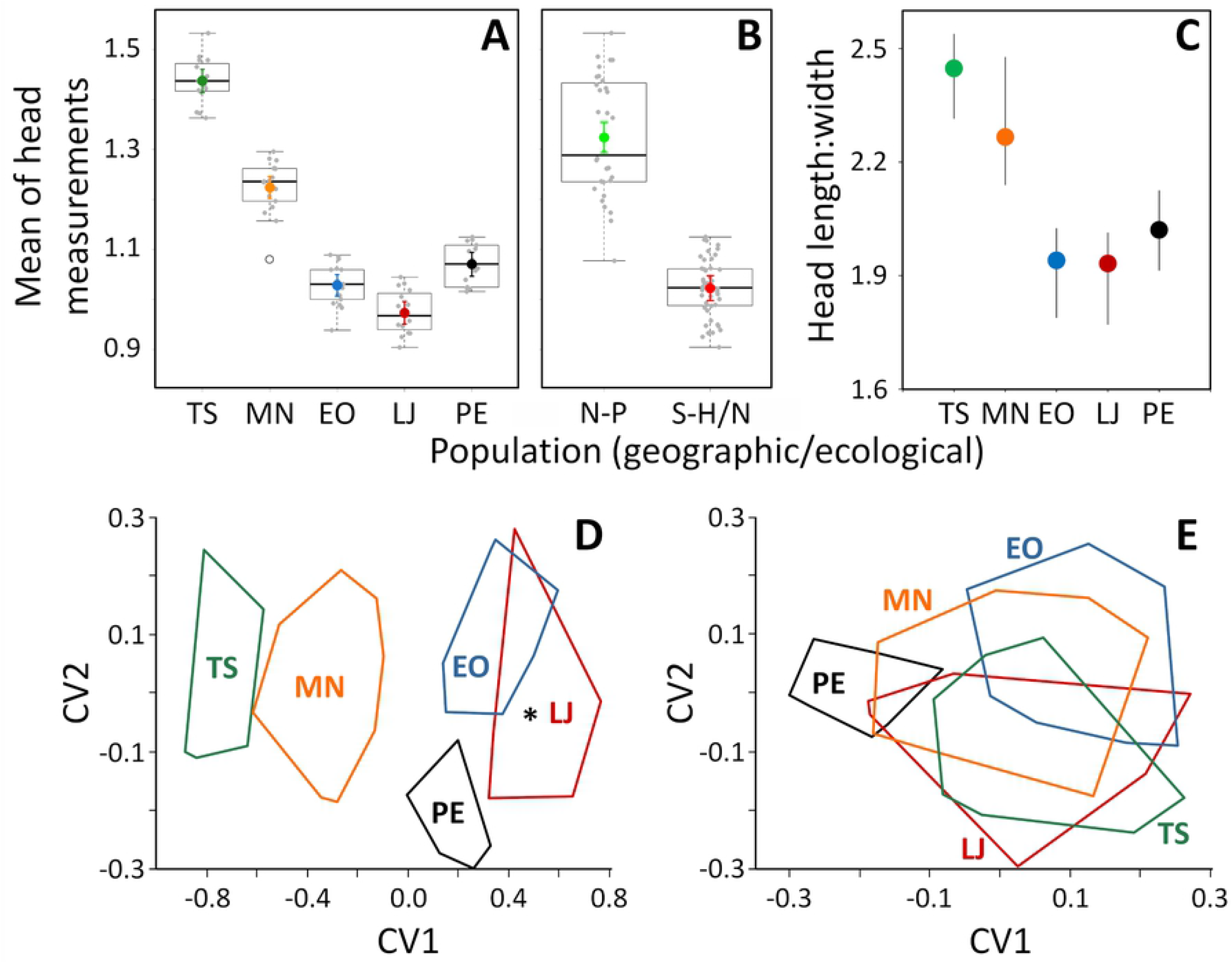
Traditional morphometrics of *Rhodnius ecuadoriensis* heads. Upper panels: head size divergence among geographic (**A**) and ecological (**B**) populations. Population codes: TS, Tsáchilas (dark green); MN, Manabí (orange); EO, El Oro (blue); LJ, Loja (dark red); and PE, Peru (black) (in **B**: N-P, northern-palm [TS+MN, light green]; S-H/N, southern house/nest [EO+LJ+PE, bright red]). Box plots show medians (thick horizontal lines), quartiles (boxes), and values that fall within 1.5 times the inter-quartile range (whiskers); note one small-sized outlier (empty circle in panel **A**) in the MN population. Colored circles and error bars show population means and 95% likelihood-profile confidence intervals. Panel **C** shows the relation between head length (yellow dotted line in Fig 2) and width (measurement ‘A’ in Fig 2), as the mean and range of raw length:width values; note the elongated heads of TS and MN bugs and the shorter-stouter heads of LJ, EO, and PE bugs. The lower panels examine size and shape divergence among geographic populations: **D**, isometry-free canonical discriminant analysis on linear head measurements; the asterisk indicates the position of a specimen from Suyo (Piura, Catamayo-Chira basin, Peru) in discriminant space; and **E**, size-free canonical discriminant analysis on the residuals of linear regression of the first principal component on each head measurement. CV, canonical vector; see text for methodological details.

**Fig 6.**
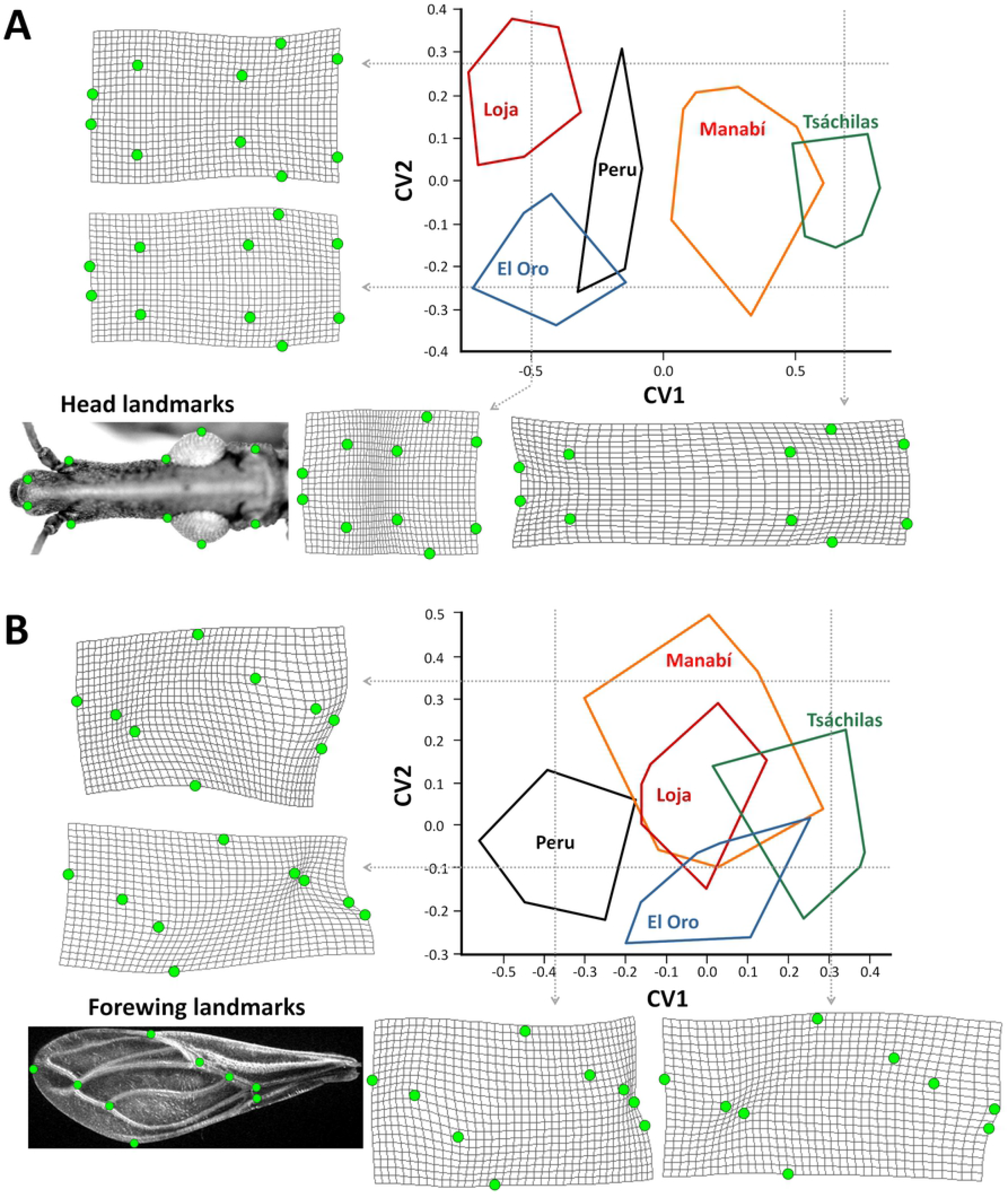
Geometric morphometrics of *Rhodnius ecuadoriensis* heads (A) and forewings (B): canonical discriminant analysis of relative warps. The plots are factorial maps based on the two first canonical vectors (CV), with convex hulls enclosing individual bugs of each population. Grids show head and forewing thin plate spline configurations at different CV values (dotted arrows); head and forewing landmarks are shown (as light-green dots) for reference. Note the striking elongation of the head with increasing CV1 scores (in **A**) and the distinctive configuration of the forewing in Peruvian bugs, with low scores on both axes (in **B**).

### Geometric morphometrics – heads and forewings

Geometric head-shape analyses (Wilks’ λ = 0.051, *p* < 0.0001) confirmed the striking contrast between the larger, elongated heads of northern palm-dwelling bugs (and, in particular, northern-Andean bugs from Tsáchilas) and the smaller, short-and-stout heads of southern-Andean house/nest bugs (Fig 6A; see also Figs 4 and 5C). The patterns revealed by CVA of forewing shape components (Fig 6B; Wilks’ λ = 0.016, *p* < 0.0001) were comparable to those revealed by size-free traditional head morphometrics (see Fig 5E), although the divergence of Peruvian bugs was clearer against a background of broad similarity among Ecuadorian populations (Fig 6B). Thin plate splines and CV1 scores suggested that most Tsáchilas, some Manabí, and a few El Oro bugs had more elongated forewings, particularly in comparison with Peruvian material; most bugs from El Oro and Loja, as well as some Manabí specimens, were somewhat intermediate (Fig 6B).

### Molecular analyses 1 – mtDNA

We found ten *cytb* haplotypes, some recovered from different collection sites, in our *R. ecuadoriensis* samples (see S1 Table, Fig 7, and below). All 663-bp sequences comprised an open reading frame with no stop codons or any other signs of pseudogene sequences. The *R. ecuadoriensis cytb* alignment had 42 variable sites (6.3%); 36 were in 3^rd^ codon positions, five in 1^st^, and one in a 2^nd^ codon position. There were two non-synonymous point mutations in a single codon of the only haplotype (PE) found in all 13 Peruvian bugs – with ACT (threonine) instead of GTT (valine). One Manabí haplotype (MN5) also had a 1^st^ codon position non-silent substitution – ATT (isoleucine) instead of CTT (leucine) (see S1 Alignment; sequences were deposited in GenBank under accession codes MT497021–MT497035 for *R. ecuadoriensis* and MT497036–MT497038 for outgroup taxa).

**Fig 7.**
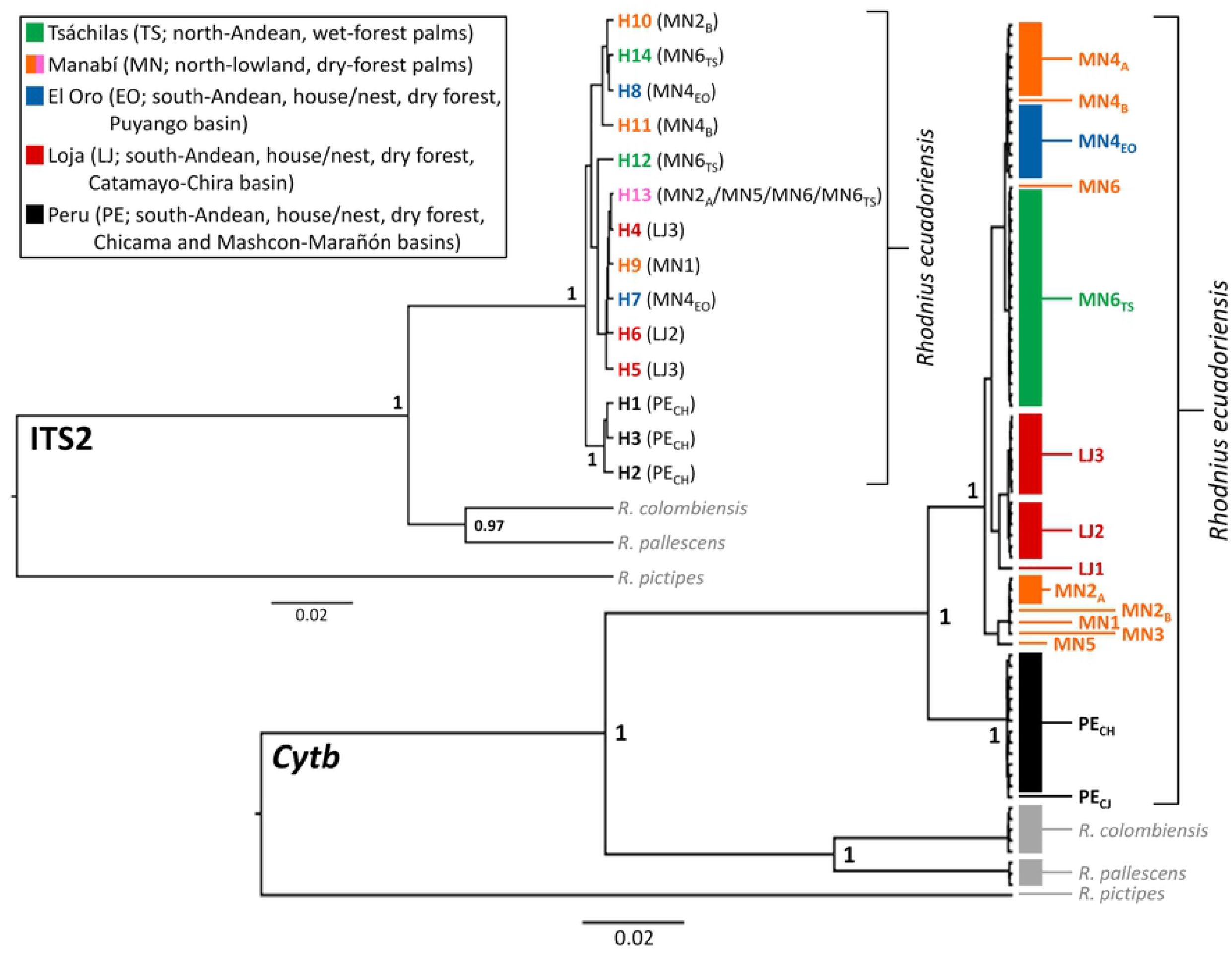
Phylogenetic relations among mitochondrial *cytb* and nuclear ITS2 haplotypes in *Rhodnius ecuadoriensis*. Mitochondrial *cytb* haplotypes found in bugs with each ITS2 haplotype are shown in parentheses in the ITS2 tree. Note the lack of differentiation among Ecuadorian populations. Note also (i) that ITS2 haplotype H13 (pink font) was found in several bugs from Manabí and in one bug from Tsáchilas and (ii) that good-quality ITS2 sequences could not be determined for bugs with *cytb* haplotypes MN3 (Manabí, Ecuador) and PE_CJ_ (Cajamarca, Peru). Bayesian posterior probabilities >0.95 are shown close to key nodes. Scale bars in substitutions/site. Outgroup taxa are in grey font.

We isolated six *cytb* haplotypes (MN1 to MN6) from northern-lowland Manabí bugs (S1 Table). MN1, MN3, and MN5 were found in one specimen each. MN2 was detected in bugs from two sites (codes MN2_A_ and MN2_B_); MN2 is identical to KC543509 from Santa Ana, Manabí [70]. MN4 was also found in bugs from two distinct Manabí sites (MN4A and MN4_B_), as well as in all nine southern-Andean bugs from El Oro (MN4_EO_) including those collected in two different sites and in different years; MN4 is identical to KC543510, also from Santa Ana, Manabí [70]. MN6 was found in one bug from Manabí and in the 20 northern-Andean specimens from Tsáchilas palms (MN6_TS_). MN4 and MN6 differ by a single, 3^rd^ codon position C/T transition. We isolated three unique haplotypes (LJ1, LJ2, and LJ3) from the 15 southern-Andean Loja bugs (S1 Table); LJ2 is identical to KC543508 from Quilanga, Loja, in the Catamayo-Chira basin [70]. Finally, 13 Peruvian bugs from at least three dwellings of the middle-upper Chicama basin also yielded a single, unique haplotype (PE_CH_). One specimen from the reference *R. ecuadoriensis* colony at LNIRTT, founded in 1979 with bugs from Cajamarca, had the same haplotype (coded PE_CJ_) (S1 Table). Haplotype PE was the most distinct among all the *R. ecuadoriensis cytb* sequences we studied; it was separated by 25 nucleotide substitutions from the closest Ecuadorian haplotype (MN1; see S2 Table and below).

Overall, uncorrected *cytb* nucleotide diversity was π = 0.0179±0.0029 SE (SEs estimated with 1000 bootstrap pseudo-replicates). Mean Kimura 2-parameter (K2p) distances were 0.0185±0.0031 SE for the 72-sequence *cytb* alignment and 0.0184±0.0029 SE for the 10-haplotype dataset. K2p sequence divergence was substantially lower among Ecuadorian haplotypes (0.0015 to 0.01995) than between any of these and PE (0.03904 to 0.04894) (S2 and S3 Tables).

The best-fitting model for the *cytb* alignment had three substitution rates (AT=CG=GT; AG=CT; and AC) with a Gamma shape parameter (+Γ) and a proportion of invariable sites (+I). The *cytb* gene tree shows an unambiguous separation (Bayesian posterior probability BPP = 1.0) of the Peruvian haplotype from a monophyletic (BPP = 1.0) Ecuadorian clade. No clear patterns of geographic or ecological segregation are apparent within the Ecuadorian clade, although the three LJ haplotypes unique to Loja bugs (see Fig 7) cluster together with BPP ≈ 0.95.

### Molecular analyses 2 – nDNA

We identified 14 unique ITS2 haplotypes (GenBank codes KT267937–KT267950) in our *R. ecuadoriensis* sample, including three (H1 to H3) from Peruvian bugs carrying the PE_CH_ *cytb* haplotype and 11 from Ecuadorian bugs – two from northern-Andean bugs (Tsáchilas, H12 and H14); three from northern-lowland Manabí bugs (H9 to H11); one (H13) from several Tsáchilas and Manabí bugs; two from El Oro (H7, H8); and three from Loja (H4 to H6) (see S1 Table). All these ITS2 sequences were overall similar to each other (see S4 Table), and no chromatogram had signs of pseudogene sequences. The 720-bp long *R. ecuadoriensis* 14-haplotype alignment (S2 Alignment) had 18 variable sites (2.5%) and 10 indels (from one to five positions long; 16 single-position gaps overall). Within Ecuador, haplotypes H4 (Loja) and H13 (Manabí and Tsáchilas) differed by a single-nucleotide indel. Using the ‘pairwise deletion’ option in MEGA X [57], we found an overall, uncorrected nucleotide diversity π = 0.0052±0.0014 SE; the values were π = 0.0046±0.0014 SE for Ecuadorian sequences and π = 0.0047±0.0021 SE for Peruvian haplotypes, with a mean uncorrected between-group distance of 0.0062±0.0019 SE.

For the ingroup + outgroup alignment (S3 Alignment; outgroup sequences with GenBank codes KT351069–KT351071), the smallest-BIC model of nucleotide substitution included four rates (AC=GT; AG=CT; AT; and CG), a Gamma shape parameter (+Γ), and a proportion of invariable sites (+I). Phylogenetic analysis revealed no geographic or ecological structuring among Ecuadorian bugs, but the three ITS2 haplotypes from Peruvian specimens clustered in a separate clade with BPP = 1.0 (Fig 7). This lent support to our *cytb* findings and, importantly, indicated that the similarity of mtDNA sequences across phenotypically and ecologically distinct Ecuadorian bugs (including the highly divergent Tsáchilas specimens) is not due to introgression [74].

### Molecular analyses 3 – species trees and Bayesian hypothesis testing

Table 2 summarizes the results of our assessment of the competing hypotheses about the number of independent lineages (one *vs*. two) within *R. ecuadoriensis*. We found that the ‘two-lineage’ hypothesis, H_1_, has very strong support from our DNA-sequence data, with log-mL estimates consistently larger (by >24 and >12 units, depending on the estimation procedure) than those of H_0_ (Table 2). These nested- and path-sampling Bayes factor estimates correspond to Bayesian posterior probabilities between 0.999998 and 1.0 in favor of H_1_; conversely, then, we found that our data provide virtually no support for the ‘single-lineage’ hypothesis (Table 2). Multispecies-coalescent analyses, therefore, substantiated locus-specific and size-free morphometric findings in that the best-supported species tree corresponds to the ‘two-lineage’ hypothesis. This tree (Fig 8) shows a well-supported *R. ecuadoriensis* clade within which our study specimens consistently segregate into two closely-related lineages: (i) the Ecuadorian lineage including typical southern-Andean house/nest bugs, atypical northern-Andean palm bugs, and intermediate northern-lowland palm bugs, and (ii) the Peruvian lineage including southern-Andean house/nest bugs (phenotypically similar to type material) from the Chicama and Mashcon-Marañón basins (Fig 8 and Figs 1, 3–6).

**Table 2.**
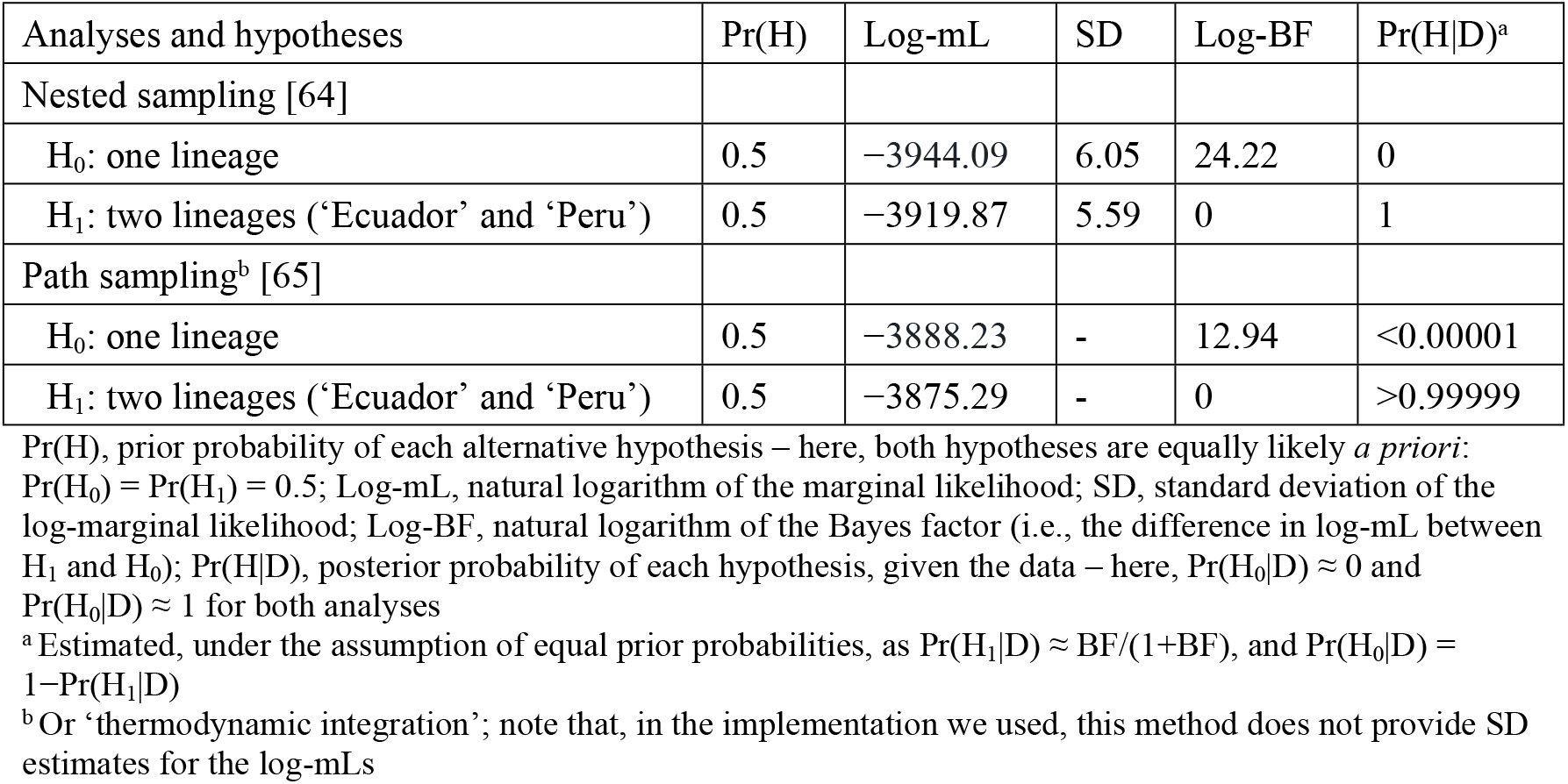
Marginal likelihoods, Bayes factors, and hypothesis testing: one *versus* two independently-evolving lineages in *Rhodnius ecuadoriensis*.

**Fig 8.**
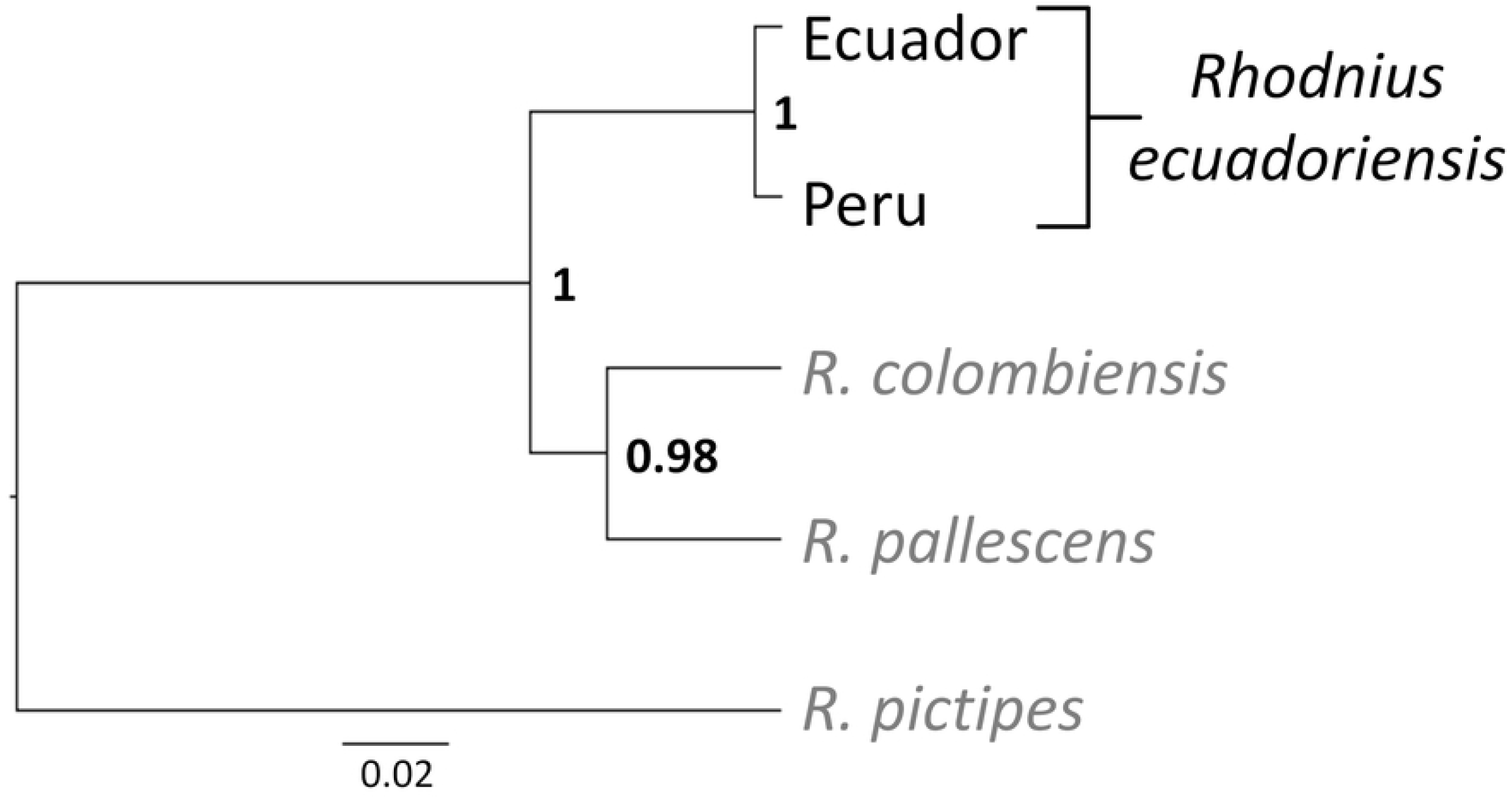
Multispecies coalescent analysis results: *Rhodnius ecuadoriensis* species tree estimated using mitochondrial *cytb* and nuclear ribosomal ITS2 sequences under the two-lineage hypothesis (‘Ecuador’ and ‘Peru’). Maximum clade credibility tree based on 3000 replicate trees; Bayesian posterior probabilities for cladogenetic events are given close to each node. Scale bar in substitutions/site. Outgroup taxa are in grey font.

## Discussion

In this report we describe a striking instance of phenotypic divergence and convergence within a single nominal species of Triatominae, *Rhodnius ecuadoriensis*. We found (i) sharp, naked-eye phenotypic divergence of genetically similar Ecuadorian bugs (southern-Andean house/nest *vs*. northern palm populations – and, within northern palm-dwelling bugs, Andean *vs*. lowland populations); and (ii) marked, naked-eye phenotypic similarity, most likely due to convergence, of southern-Andean house/nest populations (Peru *vs*. southwestern Ecuador) whose distinct DNA sequences and forewing (plus, to a lesser extent, head) shapes strongly suggest incipient evolutionary divergence (Fig 9 and S2 Fig). Below we argue that local adaptation to distinct microhabitats is probably the key driver underpinning this remarkable example of population-level phenotypic diversity.

**Fig 9.**
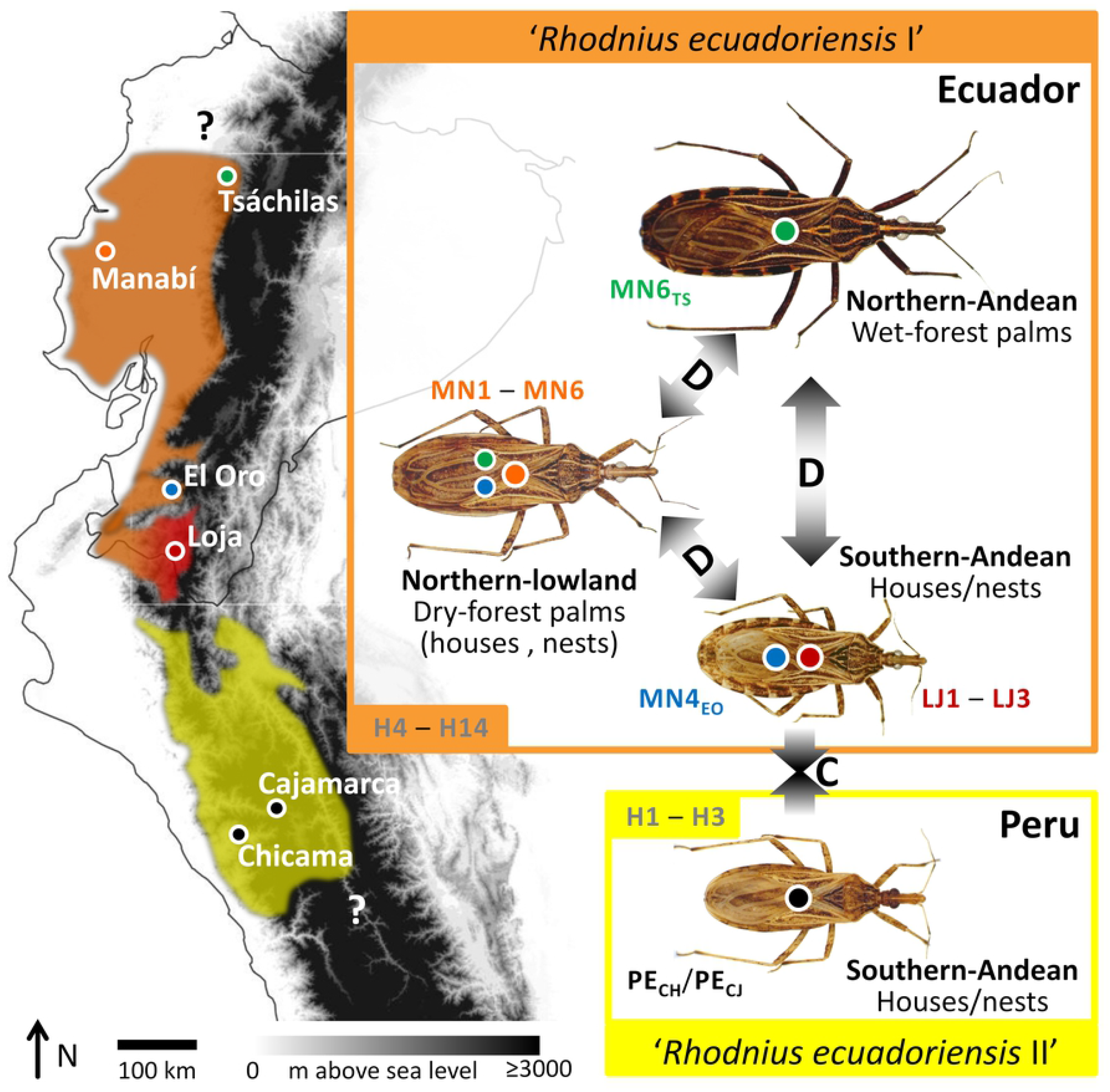
Divergence and convergence in Triatominae: genotypes, phenotypes, and habitats of *Rhodnius ecuadoriensis* populations. The map illustrates the approximate distribution of the two *R. ecuadoriensis* lineages: the Ecuadorian lineage (orange; *‘Rhodnius ecuadoriensis* I’ of refs. [8,18]) and the Peruvian lineage (yellow; ‘*Rhodnius ecuadoriensis* II’ of refs. [8,18]). Question marks highlight uncertainties as to the species’ northern and southern range limits. The reddish shade in Loja suggests possible, partial differentiation of local populations in the Catamayo-Chira basin, as indicated by the identification of three closely-related *cytb* haplotypes (LJ1 to LJ3) not shared with other populations (Fig 7) and by limited microsatellite [67] and 2b-RAD genotyping data [75]. Colored dots show the approximate geographic location (on the map) of *cytb* haplotypes and their correspondence with each phenotype (on bug pictures); color codes are as in Figs 1 and 5–7. Nuclear ITS2 haplotypes differ between Ecuadorian (H2 to H14; orange box) and Peruvian bugs (H1 to H3; yellow box), with no clear geographic, ecological, or phenotype-related genetic structuring within Ecuador. Grey-white arrows emphasize phenotypic divergence (D) or convergence (C) between populations.

Triatomines are blood-sucking bugs that live in sheltered microhabitats with a more-or-less stable food supply [1,2,27]. All *Rhodnius* species, for example, are primarily arboreal; most are tightly associated with palm-crown habitats, but some species and populations also exploit vertebrate nests built on tree branches, inside tree hollows, in bromeliads, or on palm crowns [2,27,29]. Populations of a few *Rhodnius* species have also adapted to man-made habitats and can transmit *T. cruzi* to people and their domestic mammals [1,3,27]. *Rhodnius ecuadoriensis* is one such species. In the wild, it seems to be primarily associated with the endemic *Phytelephas aequatorialis* palm of western Ecuador, but has also been found in vertebrate nests; southern-Andean populations, in particular, occur in dry ecoregions in which palms are rare or absent, and seem to have shifted to squirrel, bird, and opossum nests [1,2,26–34]. In addition, some *R. ecuadoriensis* populations can infest houses and peridomestic structures – with a preference for hen nests, guinea-pig pens, or dovecotes [1,24–27,35,36,38,39]. These synanthropic populations are important local vectors of human Chagas disease [1,24–27,36–39]. At a broader spatial scale, *R. ecuadoriensis* is the only *Rhodnius* species known to occur on the western side of the Andes south of the Magdalena-Urabá moist forests of northwestern Colombia; the Chocó rainforests along the Colombian Pacific coast separate *R. ecuadoriensis* from its sister-species clade, *R. pallescens*–*R. colombiensis* [8,18]. Within its range, *R. ecuadoriensis* occurs in widely different ecoregions [8,18,25,26]. In central-western Ecuador, presence records range from Andean wet premontane (or ‘cloud’) forests to semiarid parts of the coastal lowlands [25]. In southwestern Ecuador and northwestern Peru, the species occupies seasonally dry inter-Andean valleys up to 2700 m above sea level, and is often found infesting houses [25,26].

We reasoned that the broad ecological flexibility of *R. ecuadoriensis* was likely to correlate with similarly broad intraspecific variation, and set to examine the signs of diversification and adaptation in this locally important vector species. To address both macro-scale diversity and micro-scale adaptations, we analyzed mitochondrial and nuclear DNA markers that have proven useful in similar study-systems [5–9,11,12,15–17,19,67–70,76–80] and undertook a detailed qualitative/quantitative phenotypic assessment including head and forewing morphometrics [22,45,47–49,81]; we then used rich, specimen-specific ecological metadata to guide the interpretation of results. We nevertheless note that our results are based on a relatively limited (albeit overall well representative) sample of geographic-ecological populations and on just two genetic loci (albeit two that are known to be informative for problems similar to the one we tackled). Our interpretations of these results, therefore, are best viewed as testable hypotheses to be addressed by future research based on wider population, specimen, and character sampling.

### Macro-scale diversity: lineages and shape patterns

Our results provide strong support to the view that *R. ecuadoriensis* is composed of two major, independently-evolving lineages [8,18]. The primarily Ecuadorian lineage has been dubbed ‘*R. ecuadoriensis* group I’ [18]; it occupies highly diverse ecoregions from the wet central-western Ecuadorian Andes down to the drier valleys of the Catamayo-Chira basin – apparently always north of the Sechura desert-Huamaní range (Figs 1 and 9). The Peruvian lineage, or ‘*R. ecuadoriensis* group II’ [18], occurs in the dry inter-Andean valleys of northwestern Peru, from the Huancabamba depression down to (and apparently excluding) the semiarid Santa river basin [8,25,26]; this distribution includes (i) Pacific-slope valleys, from the eastern edge of the Sechura desert to the Chicama and perhaps Moche basins, and (ii) the Amazon-slope upper Marañón valley (Figs 1 and 9). *Cytb* divergence levels suggest [17,79] that these two lineages may have been evolving independently for 2.2–3.6 million years, with a late Pliocene-early Pleistocene most recent common ancestor. K2p distances (4.0–4.9%) are larger than those separating *R. prolixus* from its sister species, the partly sympatric *R. robustus* I (3.0–3.3%) [8,17].

Although these differences are in the limit of what Wiemers and Fiedler [82] consider ‘low’ levels of mtDNA K2p sequence divergence between reciprocally monophyletic sister clades, our ITS2 (Fig 7) and multispecies coalescent results (Table 2 and Fig 8) lend further support to the hypothesis that Ecuadorian and Peruvian *R. ecuadoriensis* are independently-evolving lineages [8,18,62]. An allozyme electrophoresis study including *R. ecuadoriensis* colony bugs originally from Ecuador (Manabí and El Oro) and Peru (Cajamarca; same colony as PE_CJ_) provides additional insight on divergence at multiple nuclear loci; in particular, different alleles of *Mdh, Pep3*, and *Pep4*, for which no heterozygotes were detected, segregated “according to the geographical origin of the specimens from Ecuador and Peru” (ref. [83], p. 303). A later study showed that the 45S rDNA gene cluster is located in different chromosomal loci in Ecuadorian (X and Y chromosomes; bugs from Manabí) and Peruvian specimens (only in X chromosomes; bugs from La Libertad) [84]. Taken together, our DNA results and these independent findings are strongly suggestive of a relatively long history of independent evolution of Ecuadorian and Peruvian *R. ecuadoriensis* lineages, likely involving ongoing or very recent speciation. Our detailed appraisal of phenotypes (Table 1, Figs 3, 4, and S1 Fig; see also S1 Text) provides the basis for distinguishing bugs carrying Peruvian and Ecuadorian genotypes; we expect professional taxonomists to examine our findings and, if warranted, formally describe a new *Rhodnius* species based on Peruvian material. If confirmed, this would have practical implications, because Chagas disease control programs could be designed to target each lineage/species separately.

Our DNA-based results, in addition, broadly mirrored those of forewing and (size-free) head shape analyses; as previously suggested [47,49], explicit size partitioning was necessary to single out genetically distinct groups after traditional morphometrics. We expected wing shape to be relatively conserved because of the crucial role of flight in adult-bug dispersal [1,27] and the importance of wing geometry for flight efficiency [85]. Forewing shape differences reflected the relatively deep genetic divergence of the Ecuadorian and Peruvian lineages (Figs 6B and 7). On the other hand, the adaptive value of head-shape variants within triatomine-bug species remains obscure. Our results suggest that, when size effects are removed, *R. ecuadoriensis* head shape may fairly mirror genetic divergence (Figs 5E and 7). In general, the elongated heads of palm-dwelling populations (Tsáchilas and Manabí) sharply contrast with the shorter, stouter heads of southern-Andean populations (Figs 3, 5C, and 6A). As discussed below, this may be related to a transition from palm to nest microhabitats [2,86].

### Phenotypic variability at the micro-scale: microhabitat adaptations

Elongated heads and medium-sized bodies (relative to other triatomines) are typical of the genus *Rhodnius*, which is mainly comprised of palm-living species [1,2,29]. The few exceptions to this rule seem to correspond to nest-dwelling species [2]. The *Psammolestes* are an atypical *Rhodnius* sub-lineage [5,6,8–10,13] with strikingly distinct phenotypes – very small bodies and very short-and-stout heads [1,2]. These clearly derived traits are probably related to the adaptation of the *Psammolestes* common ancestor to the enclosed vegetative nests of some ovenbirds [2,18,22]. Among *Rhodnius* species, the most similar in head shape and body size to typical, southern forms of *R. ecuadoriensis* is *R. paraensis*, which has so far only been reported from the tree-hole nests of arboreal *Echimys* spiny rats [1,2,27,87]. *Rhodnius domesticus* also has a relatively short and stout head for the genus; it is too among the few *Rhodnius* species not specializing in palm habitats – instead, it is associated with the nests and shelters of *Phillomys* tree-rats and *Didelphis* and *Marmosa* opossums in bromeliads and hollow tree-trunks [1,2,27].

These observations suggest that the overall reduced body size and head dimensions of typical *R. ecuadoriensis* populations may be a consequence of their shifting from the original palm-crown habitat to new, squirrel and/or bird nest microhabitats [2]. This shift likely occurred in the dry Andean environments of southwestern Ecuador-northwestern Peru that overall lack native palm populations [25,26,30,32,39]. In Loja, wild *R. ecuadoriensis* often breed inside the nests of the tree-squirrel, *Sciurus stramineus/nebouxii* [2,30,32,33,88]. Within nest microhabitats, the close physical proximity between the (virtually ectoparasitc) bugs and their hosts would relax selection for long/narrow heads and mouthparts, which may be required for biting free-ranging hosts more safely (at a longer distance) and sucking their blood faster (thanks to larger cibarial-pump muscles) [86,89]. In addition, and as has been also postulated for domestic triatomine populations [4], an overall more predictable food supply within a nest (or human dwelling), with a higher likelihood of repeated smaller bloodmeals, would relax the need for growing bigger bodies capable of storing larger amounts of blood [86]. Finally, host-mediated (passive) dispersal is probably more important among nest-dwelling than among palm-dwelling bugs, which might reduce the need for highly-efficient flight, hence relaxing selection for elongated wings [27,85]. We note that in Manabí *R. ecuadoriensis* occurs both in *Ph*. *aequatorialis* palms [2,28,29,31] and in bird and mammal nests built on palms, trees, or bromeliads [2,31]. This might help explain the intermediate phenotypes, including body size and head/forewing shape, of these lowland palm populations (Figs 3, 5, and 6).

But perhaps the most evident adaptation among the characters we studied is the striking color variability among *R. ecuadoriensis* populations. In particular, the dark hue of Tsáchilas bugs differs markedly from the typical yellowish, straw-like color of the remaining populations (Fig 3). This straw-like coloration is shared by *R. pallescens* and *R. colombiensis* [1,73], suggesting that it is plesiomorphic (S2 Fig). In the bugs we studied, color variation involved mainly pigmentation intensity rather than discrete changes in the arrangement of markings. For example, some typical-color bugs may have a larger amount of irregular dark spots and markings, or their dark markings may have larger surfaces. The highly divergent Tsáchilas forms have large and abundant black markings on a reddish-brown, generally very dark background color. Although Tsáchilas and Manabí populations share *Ph. aequatorialis* as their primary ecotope, extensive field observations [28,72] led us to notice that palm-crown microhabitats are often quite different in the wet Andes and the dry lowlands. In the dry Manabí lowlands, dead palm fronds and fibers tend to dry up, resulting in a straw-colored habitat substrate. In contrast, dead palm fronds and fibers quickly decay in the wet Andes foothills – where, in addition, large amounts of epiphytes grow on the palms. As a result, the palm-crown microhabitat of the northern-Andean Tsáchilas bugs has a darker, actually reddish-brown, background color. The color of palm-dwelling bugs from each area (Fig 3), hence, closely matches their palm-microhabitat background, suggesting camouflage against the substrate [87,90–92]. Also in line with this ‘camouflage hypothesis’, southern *R. ecuadoriensis* populations (Fig 3) associate with squirrel and/or bird nests made of light brown-yellowish materials – twigs, dry grass/leaves, and straw. We therefore suggest that sight-guided predators provide the main selective pressure underlying color variability in *R. ecuadoriensis* – and probably in other triatomines.

## Conclusions

Adaptation of an organism to its habitat becomes particularly evident when a human observer can predict habitat traits from organism traits. Our findings suggest that this is likely the case with *R. ecuadoriensis* populations. Thus, bug color predicts microhabitat background color, suggesting an adaptive response to selective pressure from sight-guided predators [92]. The small body size of southern-Andean bugs, together with their shorter-stouter heads and less elongated wings, predicts that wild populations preferentially exploit nest microhabitats [2,86] – a proposition for which there is some empirical evidence, including abundant squirrel-nest populations [2,30,32,33] and a strong association of domestic bugs with hen nests and guinea-pig pens [1,26,27,35,36,39]. Importantly, we have also shown that populations with extremely divergent phenotypes can share their genetic backgrounds, at least for the two loci we examined; our sequence data indicate that genetic similarity among Ecuadorian bugs is not due to mtDNA introgression [74]. In addition, our data reveal that southern-Andean *R. ecuadoriensis* populations with near-sibling naked-eye phenotypes belong in two distinct evolutionary lineages – the Ecuadorian *‘R. ecuadoriensis* I’ and the Peruvian *‘R. ecuadoriensis* II’ (*sensu* [18]). This can have implications for taxonomy and, hence, for the interpretation of taxonomy-dependent research results; we note, for example, that the reference *R. ecuadoriensis* strain kept at the LNIRTT has the PE_CJ_ *cytb* haplotype, which is nearly 5% divergent from the LJ haplotypes that geographically correspond to the species’ type material from La Toma, Catamayo, Loja [41]. Peruvian- and Ecuadorian-lineage bugs are also clearly divergent in forewing and (size-free) head shape, at several allozyme loci [83], and cytogenetically [84].

In sum, our detailed appraisal of phenotypic and genetic diversity in *R. ecuadoriensis* revealed phenotypic divergence within genetically homogeneous populations and phenotypic convergence of genetically distinct lineages likely on their way to speciation. Such remarkable, bidirectional phenotypic change within a single nominal species was apparently associated with adaptation to particular microhabitats. These findings shed new light on the origins of phenotypic diversity in the Triatominae, warn against excess reliance on phenotype-based triatomine-bug systematics, and confirm the Triatominae as an informative model-system for the study of phenotypic change under ecological pressure.

## Acknowledgements

We thank CA Cuba Cuba and J Jurberg for providing bugs, and the staff of the Ecuadorian Malaria Control Service for field assistance.

## Supporting information

**S1 Table. Populations, specimen details, and haplotype codes of 106 *Rhodnius ecuadoriensis* bugs used in morphometric and/or molecular analyses.**

**S2 Table. Genetic differences between mitochondrial *cytochrome b* haplotypes (663 bp) isolated from *Rhodnius ecuadoriensis* populations.** Absolute number of nucleotide differences (below diagonal) and Kimura 2-parameter distances (above diagonal).

**S3 Table. Mean genetic differences between and within *Rhodnius ecuadoriensis* populations, based on mitochondrial *cytochrome b* haplotypes (663 bp).** Absolute number of nucleotide differences (below diagonal) and Kimura 2-parameter distances between (above diagonal) and within populations (diagonal). Numbers in parentheses are standard errors calculated with 1000 bootstrap pseudo-replicates.

**S4 Table. Genetic differences between nuclear ITS2 haplotypes (H1 to H14; 720-bp long alignment) isolated from Ecuadorian and Peruvian *Rhodnius ecuadoriensis*.** Below diagonal: number of nucleotide differences (and gaps in parentheses; for example, ‘(1×4+3)’ indicates that the pairwise alignment has four 1-position gaps and one 3-position gap). Above diagonal: uncorrected *p*-distances (‘pairwise deletion’ option).

**S1 Text. Detailed descriptions of the diverse *Rhodnius ecuadoriensis* phenotypes.**

**S1 Fig. Phenotype comparisons.** Details of the connexivum of Tsáchilas forms (A) and the median process of the pygophore in Tsáchilas (B), Manabí (C), and Loja bugs (D).

**S2 Fig. Phenotype-microhabitat-phylogeny correspondences.** Multispecies coalescent species tree (as in Fig 8 of the main text), with pictures (approximately to the same scale) of adult *Rhodnius ecuadoriensis* and its closest relatives – *R. colombiensis*, *R. pallescens*, and *R. pictipes*. The distribution of phenotypes along the phylogeny suggests that the common ancestor of the diverse *R. ecuadoriensis* forms was most likely a relatively large, straw-like–colored bug. Similarly, the distribution of primary microhabitats suggests that a shift of southern-Andean populations from palm crowns (green stars) to vertebrate nests (orange circles) resulted in convergence towards the small-size, short-head/wing typical *R. ecuadoriensis* phenotype; the combined star/circle symbol indicates that northern-lowland Manabí bugs are primarily palm-dwelling but may also exploit nest microhabitats. *Rhodnius ecuadoriensis* populations: N-A, northern-Andean (Tsáchilas); N-L, northern-lowland (Manabí); S-A, southern-Andean (El Oro and Loja in Ecuador; La Libertad and Cajamarca in Peru).

**S1 Alignment. Mitochondrial *cytochrome b* haplotypes in *Rhodnius ecuadoriensis* from Ecuador and Peru, plus outgroup species (*R. colombiensis, R. pallescens, R. pictipes*).**

**S2 Alignment. Fourteen nuclear ITS2 haplotypes found in *Rhodnius ecuadoriensis* from Ecuador and Peru.**

**S3 Alignment. Nuclear ITS2 haplotypes in *Rhodnius ecuadoriensis* from Ecuador and Peru, plus outgroup species (*R. colombiensis, R. pallescens*, and *R. pictipes*).**

